# Using carrier DNA in ultra-low input library preparations for next-generation sequencing

**DOI:** 10.64898/2026.01.26.701515

**Authors:** Ping Li, Jeremy Kahsen, Karen Olsson-Francis, Stefan J. Green

**Author notes:** Address correspondence to: Stefan J. Green, Genomics and Microbiome Core Facility, Rush University Medical Center, 1750 W. Harrison St., Jelke 1255B, Chicago, IL, 60612, USA. These authors contributed equally to this work (Ping Li, Jeremy Kahsen).

## Abstract

The purpose of this study was to evaluate the use of carrier DNA (i.e., exogenous DNA spike-in) for shotgun metagenome sequencing of ultra-low levels (less than 50 picograms) of metagenomic DNA. We hypothesized that carrier DNA would improve the robustness of library preparation for samples with DNA concentrations below detection by providing a tangible amount of known DNA thereby bringing total DNA concentrations closer to recommended input ranges for metagenomic library kits. We employed adaptive PCR cycling using an iconPCR instrument (N6tec) to allow dynamic thermocycling until sufficient library for sequencing was amplified, regardless of input DNA concentration. Libraries were sequenced and mapped to reference genomes of lambda and mock community organisms, and outcome measures included total reads, on-target reads, evenness of coverage across 10 organisms within each mock community, and PCR duplication rate. We demonstrate that libraries can be prepared down to 50 femtograms of input DNA, but that there is a strong correlation between input DNA concentration and PCR duplication rate. The utility of spiking in carrier DNA is equivocal as it mildly negatively impacts the observed distribution of mock communities and serves as a loss of sequencing output. Although the loss of sequencing capacity due to carrier DNA can be partially offset by reduced loss of data from PCR duplication, carrier DNA spike-in is not recommended for routine library preparation of ultra-low input samples. Adaptive cycling allows for appropriate cycling conditions when input DNA concentrations are below detection.

## INTRODUCTION

The ability to make next-generation sequencing (NGS) libraries from ultra-low amounts of DNA (e.g., 10 pg and below^1^) is important across a wide range of fields including cleanroom monitoring, extreme environmental samples such as aerosols, skin/ocular surface swabs, among others^2,3^. In most cases, extracted DNA concentrations are below detection using fluorometric quantification, but sequence data are still desired. Prior studies with mock DNA templates diluted to sub-picogram levels have shown that direct library preparation can be preferable to whole genome amplification^3^ (WGA). However, the lack of measurable DNA can be challenging for library preparation due to uncertainty in the number of PCR cycles for library amplification. The use of carrier DNA in metagenomic sequencing of such very low biomass specimens has been explored in a limited fashion previously, including in stable isotope probing metagenomic experiments^4^ and in other low biomass systems and for Oxford Nanopore sequencing (e.g., CarrierSeq^5^). In such systems, the carrier DNA is meant to bring the sample into DNA concentration ranges recommended by library preparation kits or to enable recovery of sufficient quantities of labeled DNA for metagenomics^4^.

This study aimed to develop best strategies for ultra-low biomass shotgun metagenomics using two mechanisms, including the use of a carrier DNA spike-in of lambda DNA, and an adaptive PCR cycling instrument (iconPCR, N6tec^6^) to allow dynamic thermocycling until sufficient library was amplified. Adaptive PCR cycling, as performed by the iconPCR instrument, uses real-time fluorescence data from individual reactions to determine how many PCR cycles should be performed for each sample. In the simplest form of adaptive cycling, a fluorescence threshold is set, and each sample is cycled until achieving that threshold. This is achieved by having independent control for every well in a 96-well thermocycler. For ultra-low input samples, adaptive cycling can be beneficial by allowing samples to cycle until an appropriate amount of library has been generated, especially when input sample concentrations are not known *a priori*. This contrasts with standard library preparation protocols which have recommended cycle numbers for measurable DNA inputs.

To examine these two mechanisms, mock community DNA samples and spike-in DNA samples were sheared and mixed in various ratios. These combined DNAs were diluted across a range of very low input concentrations and used as input for library preparation, followed by amplification using an iconPCR device. Subsequently, the libraries were pooled and sequenced. Outcome measures included the number of reads, percentage of on-target mapping reads, ratio of spike-in-to-mock DNA template reads, the recovery of the expected ratio of mock organisms, and the rate of PCR duplicate generation. The results indicate that sample libraries can be made from ultra-low DNA inputs using this approach, with viable performance metrics at 500 femtogram and poor, but still detectable, performance at 50 femtogram input levels. DNA spike-ins had very modest negative impacts on the recovery of mock community profiles. Adaptive PCR cycling allowed for simultaneous amplification of libraries across four orders of magnitude, and cycle number did impact the total library yield.

## MATERIALS & METHODS

### DNA templates

Three DNA templates were used for analysis in this study, including purified Enterobacteria phage lambda DNA (Lambda DNA; Oxford Nanopore Technologies, EXP-CTL001), ZymoBIOMICS Microbial Community DNA Standard (Zymo Research, D6306), and Mycobiome Genomic DNA Mix (ATCC, MSA-1010). Prior to dilution, DNA concentrations were measured using Qubit fluorimetry (ThermoFisher Scientific). DNA samples were then sheared using the RapidShear™ 24-in-5 gDNA Shearing System (Triangle Biotechnology) following the manufacturer’s protocol. Briefly, 45 µL of DNA at 2–10 ng/µL was mixed with 5 µL of RapidShear reagent and sonicated for 6 minutes. The resulting average fragment size ranged from 450 to 550 base pairs. DNA templates were then mixed in varying proportions, including 100% Zymo mock DNA (Z), 100% Lambda DNA (L), 1:1 Lambda-Zymo DNA (L1Z1), 3:1 Lambda-Zymo DNA (L3Z1), 5:1 Lambda-Zymo DNA (L5Z1), 10:1 Lambda-Zymo DNA (L10Z1), 1:1 Lambda-ATCC Mycobiome DNA (1LA1), and no-template controls (NTCs). After combining sheared template and sheared lambda DNA, samples were diluted across multiple orders of magnitude, including 5 pg/µl, 500 fg/µl, 50 fg/µl, and 5 fg/µl total DNA. Template mixing, shearing, and dilution were performed twice, independently, in workflows conducted approximately 1 month apart. In the first workflow, dilutions were performed to achieve a minimum amount of 500 fg per reaction. In the second workflow, the lowest dilution contained 50 fg per reaction.

### Library preparation

All reactions were performed in technical triplicate using a ThruPLEX DNA-Seq Kit (Takara Bio USA, Inc., San Jose, CA; catalog #R400676) with unique dual indexing according to the manufacturer’s instructions with modifications during final PCR cycling. Maximum volumes of 10 µl of sheared DNA were employed, leading to total input DNA amounts of 50 pg, 5 pg, 500 fg, 50 fg, and no input (NTCs). 50 pg is the minimum input amount recommended by the standard protocol. Library preparation is a single tube procedure including DNA repair, stem-loop adapter blunt-end ligation, extension of templates, and cleavage of loops. The final adapter-ligated templates are amplified with unique dual indexing (UDI) primers. Library amplification reactions included 0.5 µL of 200x SYBR Green I dye per 100 µL sample (1X final working concentration), and cycling was initiated on a C1000 Touch Thermal Cycler (Bio-Rad) using the following conditions: (a) Extension and cleavage: 72 °C for 3 minutes, 85 °C for 2 minutes, and denaturation: 98 °C for 2 minutes; (b) index incorporation: four cycles of 98 °C for 20 seconds, 67 °C for 20 seconds, and 72 °C for 40 seconds, and (c) library amplification (recommended 6 to 16 additional cycles) with cycling conditions of 98 °C for 20 seconds and 72 °C for 50 seconds. This last step of library amplification was performed on the iconPCR platform (N6 tec) with real-time, fluorescence-based normalization. The selected auto-normalization method chosen was “target fluorescence”, set at 10,000 relative fluorescence units (RFU), with a maximum of 20 additional PCR cycles allowed. Standard kit recommendations from Takara indicate 15-16 cycles (library amplification stage) for 50 pg inputs. Library pools were purified using KAPA Pure Beads (Roche), quantified using a Qubit4 Fluorometer with the Qubit dsDNA HS Assay Kit (Thermo Fisher Scientific), and assessed for fragment size distribution on a TapeStation device using D1000 ScreenTape (Agilent Technologies).

### Sequencing

Libraries were pooled in equal volume for each sequencing run without further attempt at library balancing. Therefore, within each experiment, read depth was an outcome measure. Library pools were sequenced on an Illumina MiniSeq platform using a paired-end 151 × 8 × 8 × 151 run configuration with a 5% PhiX control spike-in. Raw sequences from two independent sequencing runs were deposited in the Sequence Read Archive (SRA) under the Bioproject accession PRJNA1275972. Sample processing, library preparation, and sequencing were performed in the Genomics and Microbiome Core Facility (GMCF) at Rush University Medical Center.

### Contaminant identification

Although the primary approach for analysis of sequence data was through a mapping procedure against known targets (see below, data analysis), non-targeted annotation pipelines were employed to identify microbial DNA present as reagent contaminants. Once identified, whole genome sequences for these organisms were obtained and used as references in the mapping pipeline together with expected targets (i.e., mock community genomes and lambda DNA). Briefly, taxonomic classification of sequence reads generated from reagent negative controls was conducted using the Kraken2 tool, a k-mer-based taxonomic classification system^7^. The controls were analyzed using Kraken2 with a combination of reference sequence databases, including Human, Bacteria, Viruses, Fungi, and Archaea. In addition, short read sequence taxonomic annotation was also performed using the software package MetaPhlAn3 (v4.0.1^8^). Negative control taxonomic classification was performed by the Rush Research Bioinformatics Core (RRBC) at Rush University, Chicago, IL.

### Data analysis

Raw sequence data were processed within the software package CLC Genomics Workbench (v25.0.1; Qiagen). Data were initially trimmed to Q20 and mapped against mock community genomes as indicated by Zymo and ATCC and potential contaminant organism genomes (GenBank References: CP053915, CP097636, CP016210, CP069300, CP170561, CP171806, CP078069, CP077308, CP065037, CP050454, CP000555, CP002095, CP041150, LT629971, CP116346, CP119083, CP016022, CP050124, CP192575, CP124551, CP010836, CP166581, CP046508) using default mapping parameters (length fraction= 0.5 and similarity fraction = 0.8). Outcome measures included: number of reads after quality trimming, number of mapping reads, number of reads mapping to expected references, ratio of mock community to lambda DNA reads, and percentage of reads mapping to non-expected references (i.e., contaminants). In addition, a univariate metric (Ideal Score^9^) was calculated for each sample to evaluate how well sequencing recovered the expected distribution of microbial taxa within the mock community DNA. The Ideal Score was initially developed to produce a univariate metric for mock communities with even distribution of taxa, and later revised to accommodate mock communities with uneven distribution^10^. Briefly, the metric is derived from Bray-Curtis dissimilarity calculations^11^, and is a summation of the absolute difference of the expected relative abundance of each taxon and the observed relative abundance of the same taxon. A value of 0 is a perfect match between expectation and observation across the entire community, while a value of 200 is the maximum value and represents complete divergence between expectation and observation. Duplicate read levels were evaluated on reads mapping to Lambda DNA due to the greatest depth of coverage for this template relative to mock community genomes. Briefly, reads were mapped using default settings as described above. Subsequently, the “Remove Duplicate Mapped Reads” algorithm was employed within CLC Genomics. Percent duplicate levels were obtained for all samples containing at least 10,000 mapping reads to the lambda reference genome.

### Visualization

Raw counts and weighted mappings were imported into R^12^, wrangled using the tidyverse^13^, visualized with ggplot2^14^, and integrated into composite figures using patchwork^15^. Statistical differences between pairs were evaluated with t-tests, and group-level effects were analyzed with ANOVA, followed by Tukey’s post-hoc tests when applicable.

## RESULTS

Two different mock community DNA samples were separately mixed with different amounts of spike-in DNA (lambda), and libraries were prepared at DNA input levels from 50 pg to 50 fg using adaptive cycling for amplification In the first series of experiments, only the Zymo mock community was employed, and the minimum DNA input was 500 fg. In the second series of experiments, both Zymo and ATCC mock communities were employed, and the minimum DNA input was 50 fg. Average read depth, average reads mapping to mock community DNA, average reads mapping to lambda DNA, average adaptive PCR cycle number, and average reads on target (i.e., reads mapping to mock DNA and lambda DNA combined) are shown for experiments with Zymo mock community DNA in **Figure 1** and ATCC mock community DNA in **Figure 2A**. With decreasing input, adaptive cycling increased the average number of amplification cycles. In reactions with 50 fg (0.05 pg) DNA input, cycle number reached the maximum cycles allowed (20 cycles, with one exception of 19 cycles), and this was also associated with fewer total reads from 50 fg reactions. At 500 fg, a range of 18-19 cycles (1 exception with 17 cycles) was observed across all experiments, while 5 pg inputs were cycled from 16-17 cycles and 50 pg inputs were cycled from 12-13 cycles. Blanks were amplified with 19 (1 reaction) and 20 (5 reactions) cycles (**Figure 1**; **Figure 2A**).

**Figure 1.**
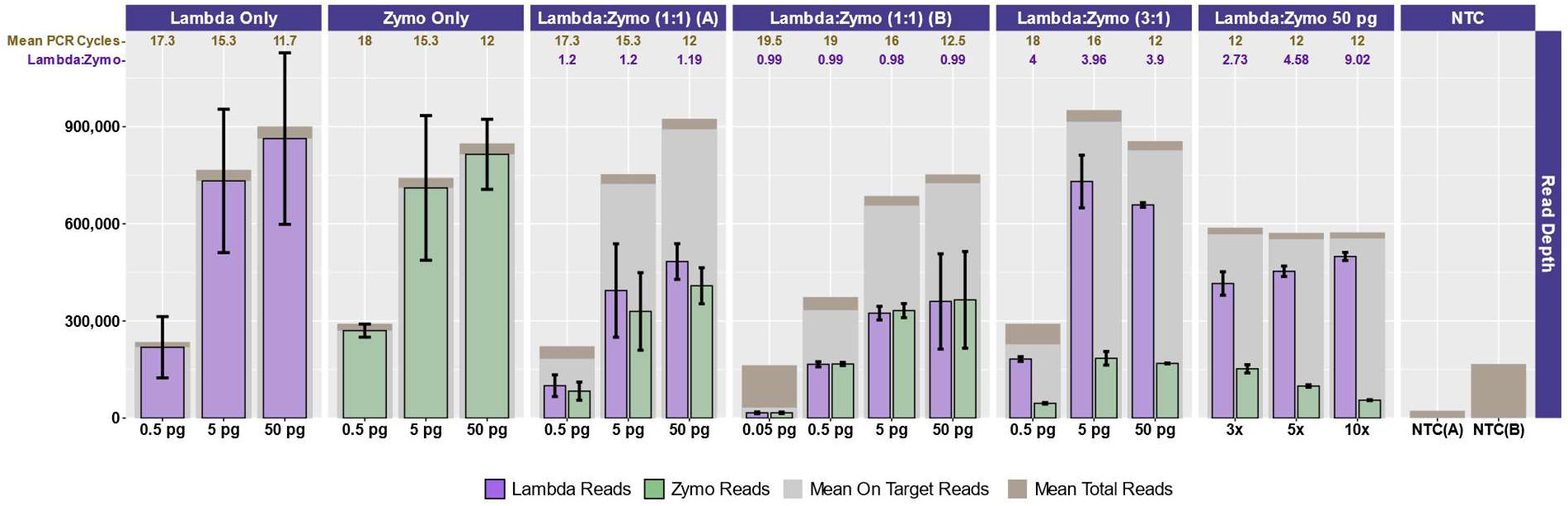
Sequencing and mapping metrics for Zymo mock community and Lambda DNA experiments. For each experiment (Lambda DNA only; Zymo DNA only; Lambda:Zymo in 1:1 input ratio – run separately twice; Lambda:Zymo in a 3:1 ratio; Lambda:Zymo in 3:1, 5:1 and 10:1 ratios; and reagent negatives from two separate runs, NTC-A and NTC-B) total read depth is shown (brown columns) together with on-target reads (*i.e.*, mapping to Lambda and Zymo genome sequences) in gray. For each experiment, the number of reads mapping to Lambda DNA (purple) and Zymo DNA (green) references are shown. Error bars indicate standard deviation associated with three replicates, except for two experiments where one replicate was dropped due to failed amplification. The average number of cycles of PCR used for amplification is shown at the top in brown numbering. The average measured mapping ratio of Lambda DNA to Zymo DNA is shown below the PCR cycle number in purple lettering. For most experiments, the input DNA amount is shown below (ranging from 0.05 to 50 pg). For the Lambda:Zymo DNA experiment comparing ratios, 50 pg inputs were used while changing the input ratio from 3:1 to 10:1.

**Figure 2.**
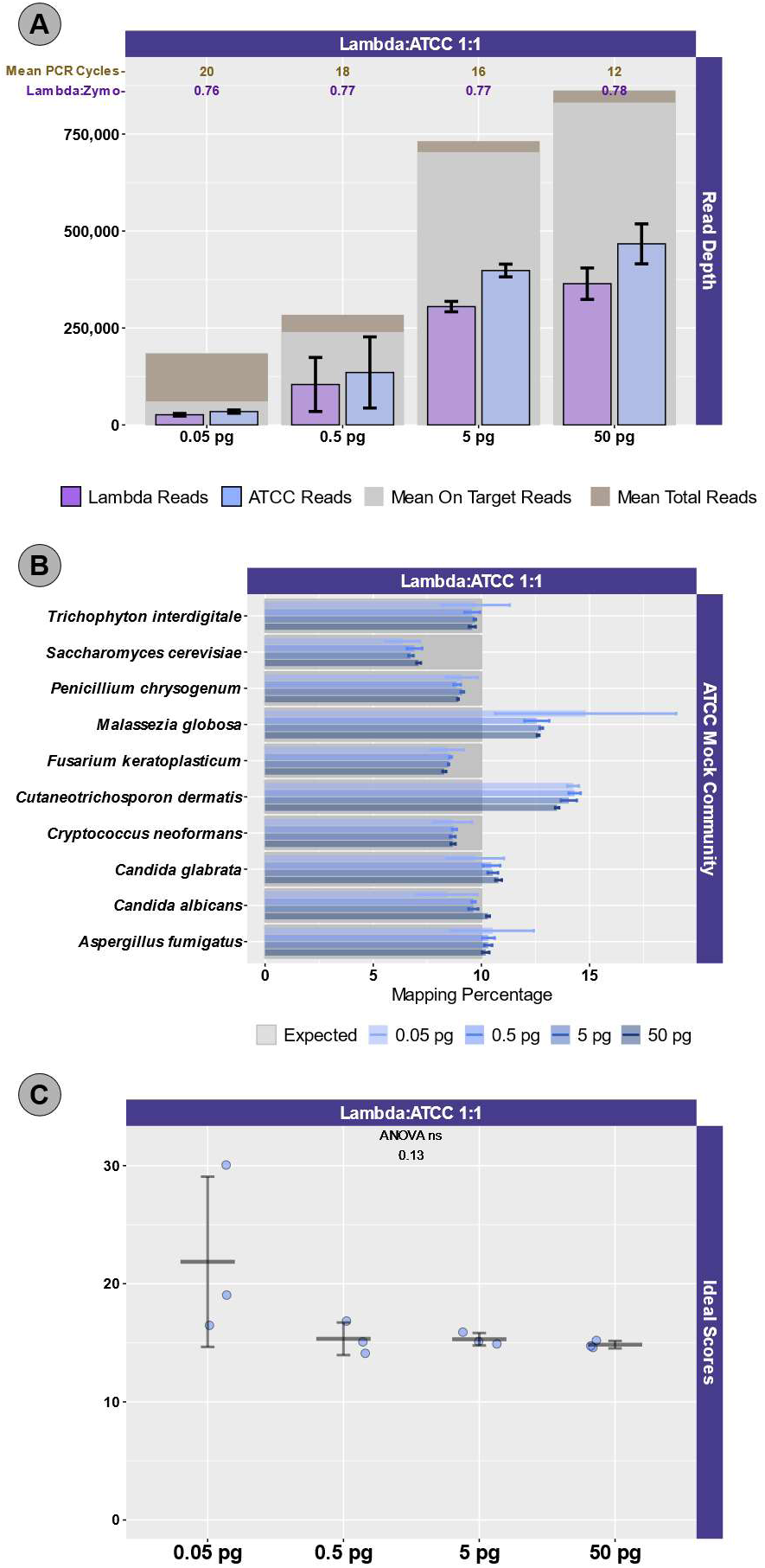
Sequencing and mapping metrics for ATCC mock community and Lambda DNA experiment. Lambda DNA and ATCC mock community DNA were sheared, mixed in a 1:1 ratio, and diluted to yield 0.05 pg to 50 pg inputs. (**A**) For each input level, total read depth is shown (brown columns) together with on-target reads (mapping to Lambda and ATCC genome sequences) in gray. The number of reads mapping to Lambda DNA (purple) and ATCC DNA references (blue) are shown. Error bars indicate standard deviation associated with three replicates. The average number of cycles of PCR used for amplification is shown at the top in brown numbering. The average measured mapping ratio of Lambda DNA to ATCC DNA is shown below the PCR cycle number in purple lettering. (**B**) The relative abundance of reads mapping to each of the 10 ATCC mock community fungal genomes are shown. Results are shown for libraries with 0.05 to 50 pg DNA input amounts. The dark gray boxes indicate the expected relative abundance of the input DNA (i.e., 10% relative abundance for each organism). Error bars indicate the standard deviation associated with three technical replicates. (**C**) Distribution of ‘Ideal Scores’ for ATCC mock community sequences across experiments. Ideal scores are a univariate metric measuring the summed divergence of observed microbial communities from expected microbial communities. The metric ranges from 0 (perfect fit) to 200, with lower numbers indicating better matching between observed and expected relative abundances across all taxa. An ANOVA was performed to determine if input DNA concentrations affected the observed microbial community structure. Error bars indicate the standard deviation associated with three replicates.

Total sequencing depth was fairly consistent for samples with 5-50 pg inputs (534,986 to ∼1,095,396 reads/sample with a mean of 767,600 reads), while sequencing depth decreased substantially at 500 fg (113,586 to 482,702 reads/sample with a mean of 282,981 reads) and 50 fg (126,844 to 216,212 reads/sample with a mean of 176,198 reads), despite greater PCR cycle numbers. Rates of on-target mapping were highest at 5 and 50 pg inputs (range of 95.28% to 96.77%; mean of 96.19%), slightly lower at 500 fg inputs (range of 75.43% to 93.63%; mean of 86.61%), and lowest at 50 fp inputs (range of 19.66% to 42.64%; mean of 28.45%) (**Figures 1 and 2A**). Negative controls (i.e., library reaction blanks) were sequenced as well, generating from 15,216 to 31,930 reads/sample (Mean of 22,683) in the first experiment and from 98,074 to 256,896 reads/sample (mean of 166,935) reads in the second experiment (**Figure 1**).

We evaluated the ratio of lambda (spike-in) to mock community sequence data against expected yields based on input DNA mixtures. In the first experiment, Lambda:Zymo DNA ratios were somewhat elevated relative to the expected ratio (**Figure 1**). For example, in dilution series from 50 pg to 500 fg, Lambda:Zymo DNAs were mixed in equimolar ratio while sequence output demonstrated elevated Lambda:Zymo (∼1.2X compared to expected 1X) (**Figure 1, Experiment ‘A’**). Similarly, for dilution series in the first experiment with Lambda:Zymo mixtures with 3X more Lambda DNA relative to Zymo DNA, sequence output ratios were elevated (∼3.9-4X compared to expected 3X). Conversely, in the second experiment (**Figure 1**, **‘Experiment B’**), the situation was reversed, with Lambda:Zymo mixtures slightly lower than expected (∼0.97-1.0X compared to expected 1X). In the second experiment, when adding Lambda DNA at increasing levels (i.e., 3X, 5X and 10X higher than Zymo), sequence yields returned slightly lower than expected ratios (2.73X, 4.58X, and 9.02X, respectively). Similarly, when Lambda DNA was mixed with the ATCC Mycobiome mock DNA at 1:1 ratios, sequence yields returned slightly lower than expected ratios (0.75-0.78X compared to expected 1X) across all input concentrations from 50 fg to 50 pg (**Figure 1**).

We next examined the distribution of reads across the expected 10 microorganisms in the ATCC **(Figure 2B**) and Zymo (**Figure 3**) standards. For the ATCC standards, highly similar profiles were observed across all experiments, regardless of input DNA concentrations or ratio of lambda DNA **(Figure 2B**). Fungal DNA sequences from the ATCC were detected in ratios close to even across 10 taxa, though two fungal taxa were somewhat enriched, including *Malassezia globosa* and *Cutaneotrichosporon dermatis* (**Figure 2B, 2C**). Slightly worse performance in recovering the expected 10% output per organism was observed at the 50 fg input level, particularly with respect to *Malassezia globose* (**Figure 2B**). In experiments with Zymo standards, very poor recovery (on average, ∼1/10^th^ of expected output; **Figure 3**) of the two fungal taxa was observed (*i.e.*, *Saccharomyces cerevisiae* and *Cryptococcus neoformans*), regardless of condition. Conversely, substantially higher than expected reads mapped to *Lactobacillus fermentum* (∼2.5X of expected output; **Figure 3**), regardless of condition. Reads mapping to *E. coli* were slightly affected by the ratio of Lambda DNA, with the highest (and greater than expected) mapping reads observed with the highest (10X higher) Lambda:Zymo ratios (**Figure 3**). Negative controls mapped poorly to the reference genomes (range of 40 to 151 reads/sample; mean of ∼81 reads), with most reads mapping to the *E. coli* reference).

**Figure 3:**
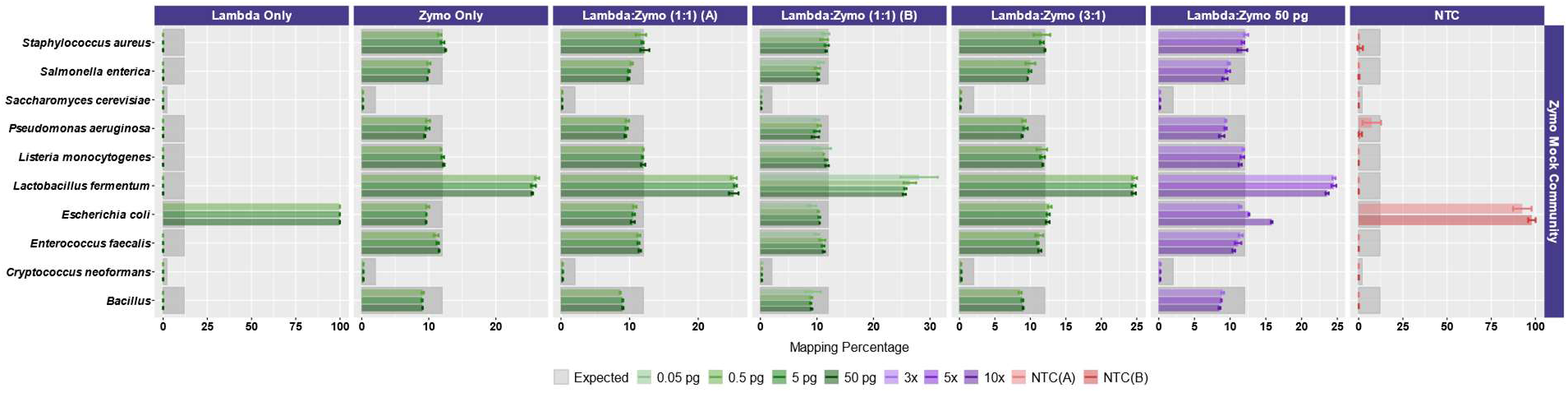
Mapping metrics for Zymo mock community sequences across experiments. The relative abundance of reads mapping to each of the 10 mock community microbial genomes are shown. The dark gray boxes indicate the expected relative abundance of the input DNA (i.e., 12% relative abundance for each of eight bacterial strains and 2% relative abundance for two fungal strains). Error bars indicate the standard deviation associated with three technical replicates (most experiments). Control data, include Lambda DNA only and no-template controls (NTC), contain relatively few reads mapping to Zymo references, and with most reads mapping to the Zymo *E. coli* reference. The x-axis represents the relative abundance of reads mapping to each of the 10 reference genomes, with the scale varying for each experiment.

To more easily compare between conditions and mock DNAs, we used a univariate metric, the ideal score, to calculate a distance between the observed distribution of reads across the 10 reference organisms and the expected distribution of reads, based on DNA ratios provided by Zymo (**Figure 4**) and ATCC **(Figure 2C**). No significant differences in Ideal scores were observed as a result of different input DNA concentrations, including experiments with 50 fg inputs. A slight increase in Ideal score, but not significantly different, was observed for the 50 fg input of a sample with a 1:1 Lambda:ATCC input DNA relative to 500 fg, 5 pg, and 50 pg samples **(Figure 2C**). Samples containing Zymo DNA had Ideal scores ranging from 25.02 to 36.81 (mean=27.61) Samples containing ATCC DNA had Ideal scores ranging from 14.11 to 30.05 (mean=16.84). Ideal scores for Zymo samples were significantly higher (P<0.001, two-tailed TTEST) relative to ATCC samples due to the poor performance of the two fungal taxa, and the overperformance of *Lactobacillus fermentum*. Negative control samples had exceedingly high Ideal scores as most mapping reads mapped to the *E. coli* reference genome from the Zymo mock community. Samples with 10:1 ratios of Lambda:Zymo had slightly but significantly higher average Ideal score relative to 5:1 and 3:1 ratio samples **(Figure 4**).

**Figure 4:**
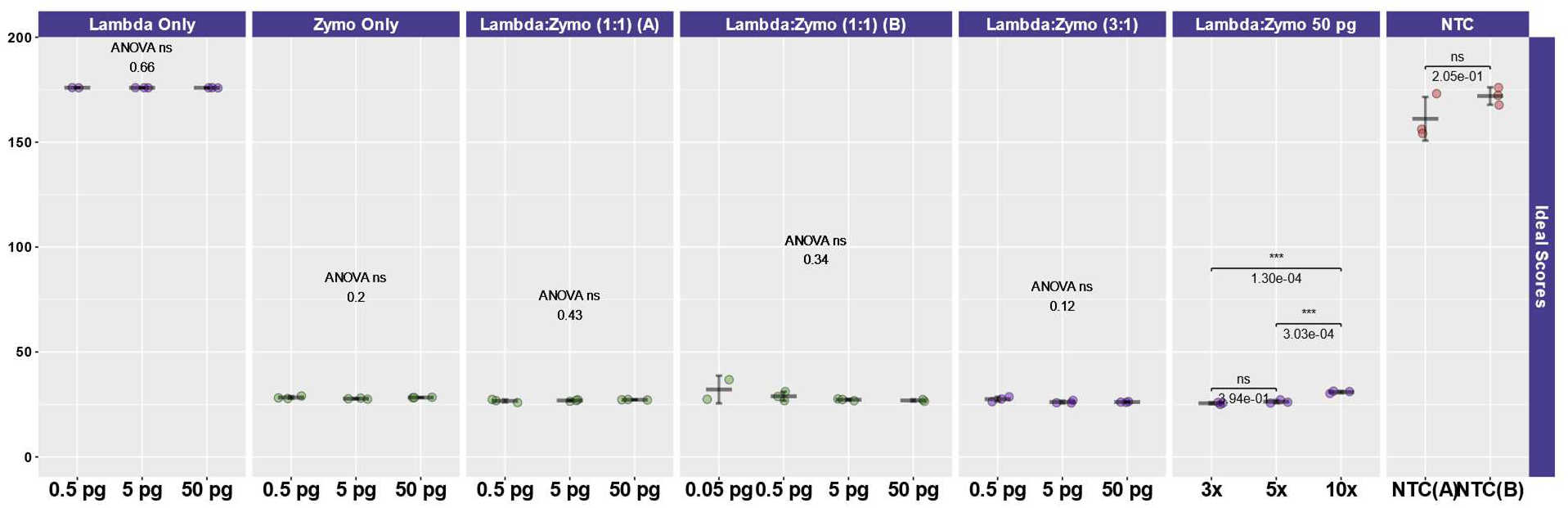
Distribution of Ideal Scores for mapping of sequences across Zymo mock community experiments. The y-axis represents Ideal scores ranging from 0 (perfect fit) to 200, with lower numbers indicating better matching between observed mapping and expected relative abundances across all taxa. Within each experiment, an ANOVA was performed to determine if input DNA concentrations affected the observed microbial community structure (most experiments) or if the ratio of Lambda:Zymo DNA affected the observed microbial community structure. Error bars indicate the standard deviation associated with three replicates (most experiments). Samples without mock community DNA (Lambda only, NTCs) have high Ideal scores due to mapping of a small number of reads to mock community genomes, particularly *E. coli*.

To evaluate the effect of input DNA concentration on library duplication, trimmed sequence reads were mapped to the lambda genome by itself. Due to the small size of the genome, deeper sequencing was generally acquired for Lambda relative to mock community DNA, which contained 10 bacterial and fungal genomes (Zymo) or 10 fungal genomes (ATCC). Thus, the Lambda genome re-sequencing was more suitable for duplication examination. **Figure 5** demonstrates the relationship between input genomic DNA and duplication rate, with a clear negative correlation between input DNA amount and duplication rate. At the highest input levels (50 pg), duplication rates ranged from a minimum of 2.92% to a maximum of 5.52% (mean=4.10%). Conversely, at the lowest input levels (50 fg), duplication rates ranged from a minimum of 75.87% to 82.68% (mean=80.26%). Duplication rate by input DNA concentration was modeled logarithmically (Duplication rate = −27.1 * log(DNA concentration, pg) + 49.9), with strong support (R^2^=0.94).

**Figure 5:**
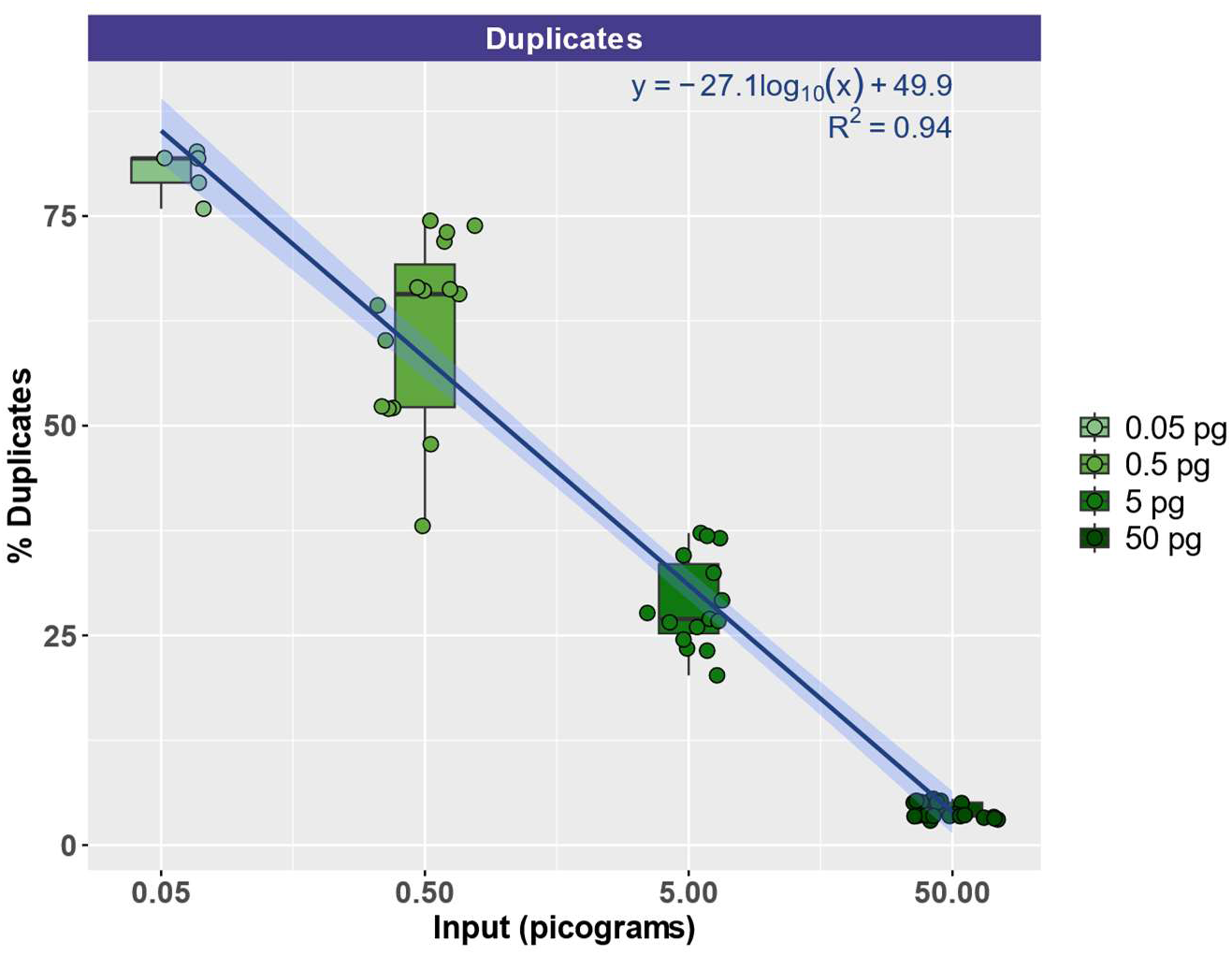
Correlation of input DNA levels with PCR duplication rates of reads mapping to the Lambda genome. A strong negative correlation between DNA input levels (x-axis; pg) and PCR duplication rate (y-axis, %) was observed. This correlation was fit with a logarithmic curve where y= −27.1 * log(x) + 49.9.

## DISCUSSION

The primary objective of this study was to determine whether non-microbial carrier DNA, such as Lambda DNA, was useful for generating shotgun metagenome sequence libraries from ultra-low DNA (<50 pg) samples. We hypothesized that carrier DNA could improve the library preparation for samples with DNA concentrations below detection by providing a tangible amount of known DNA and bringing total DNA concentrations into recommended ranges of low input library preparation reagents. Secondary objectives of this study were to evaluate whether adaptive PCR cycling, as performed by the iconPCR instrument (N6tec), could be integrated into library preparation workflows for samples with DNA concentrations below detection. The results in this study demonstrate that carrier DNA is not necessary from a workflow perspective when DNA concentrations are 500 fg and above. Below 500 fg (i.e., 50 fg), sequencing performance metrics worsened substantially, leading to fewer total reads and low on-target percentages. This is unsurprising, as at 50 fg inputs, stochastic effects based on the relatively low number of target molecules become more prominent, as does the impact of reagent contaminant DNA. In addition, the reduced number of template molecules also leads to a very high duplication rate. Thus, carrier DNA, on the scale of 1 pg, might be favored for samples where DNA levels are below detection and when a guaranteed library, regardless of input, is preferred. More broadly, however, larger amounts of carrier DNA led to slightly worsening performance in terms of recovering mock community profiles, but the primary concern with carrier DNA spike-in is the reduced sample sequence output due to the sequence yield from carrier DNA. However, the loss of sample-derived sequence yield due to the presence of carrier DNA must be considered in context of the loss of unique sample-derived sequence data due to high duplication levels observed at low input levels in the absence of carrier DNA. The total yield of non-duplicated sequence data from the sample is the primary concern in any such consideration. Conversely, sequencing performance, based on mapping reads and evaluated by Ideal Score calculations, was shown to be similar across four orders of magnitude of input DNA, and was minimally affected by carrier DNA.

The adaptive cycling of the iconPCR provides an ideal mechanism to tailor amplification strategies to input template concentrations, even when these are below the detection of high-sensitivity fluorometric input DNA concentration measurements. Additional efforts should be taken to reduce the number of PCR cycles even further using the iconPCR, to decrease the negative impact of PCR duplicate production. However, a higher maximum number of allowed cycles for amplification may be desirable when extremely low inputs are possible; in our study, 20 cycles was the maximum and with the 50 fg input led to fewer total reads relative to other input amounts. Given that an average of 17-19 cycles was observed for 500 fg input, 20 cycles was likely too few for 50 fg input amounts.

Overall, the success of this protocol should be taken in context of the fact that the manufacturer’s suggested minimum input is 50 pg, leading to library preparation from input DNA levels two-to-three orders of magnitude lower. Similar success in library preparation below manufacturer recommended inputs has been previously observed^16^, though at higher inputs than described here. Elsewhere, library preparation from ultra-low inputs (sub pg) has been previously demonstrated using Illumina Nextera library preparation^17^, with levels of duplication broadly similar to those observed here. One key advantage of the workflow in this study is the use of the dynamic cycling of the iconPCR, which allows for calibrated amplification when DNA input levels are not known. Given the robustness of the overall workflow, we recommend this as a default protocol for processing ultra-low biomass specimens from spacecraft assembly facilities and other cleanrooms, aerosol samples, deep subsurface samples, and more. Furthermore, the viability of this methodology for ultra-low DNA inputs may obviate the need for whole genome amplification (WGA) of ultra-low biomass samples, which can lead to substantial distortion or amplification bias of the underlying microbial community^18,19^. Further work will examine the use of iconPCR and carrier DNA in long-read sequencing applications, such as Oxford Nanopore^20^, where higher minimum DNA inputs are required. Limitations of the study include the use of mapping to identify on-target sequences, an approach which will not be possible for environmental samples.

## ACKNOWLEDGMENTS

The authors greatly appreciate discussions with Scott Tighe and Kasthuri Venkateswaran and others at a Planetary Protection workshop at the NASA Ames Research Center in Moffett Field, CA (November 2024), as well as comments and discussion from members of the NASA Planetary Protection Analysis Working Group. Discussions with Steve Butcher (Takara), Ioanna Andreou (Qiagen), Samuel J. Rulli (Qiagen) and Dominic ONeil (Qiagen) were also helpful. Some reagents were kindly donated by Zymo Research. This research was funded in part by a Mars Exploration Program (MEP) grant under task 108575-13.05.01.2012 carried out at the Jet Propulsion Laboratory, California Institute of Technology, under a contract with National Aeronautics and Space Administration. This research was also funded in part by a 2024 NASA Planetary Protection Research ROSES award (80NSSC24K1354) to Kasthuri Venkateswaran. Bioinformatics analysis of negative controls was kindly performed by Dr. Ankur Naqib in the Rush Research Bioinformatics Core (RRBC) at Rush University Medical Center.

## REFERENCES

1. Simpson AC, Tighe S, Wong S, et al. Analysis of Microbiomes from Ultra-Low Biomass Surfaces Using Novel Surface Sampling and Nanopore Sequencing. J Biomol Tech 2023;34(3) doi: 10.7171/3fc1f5fe.bac4a5b3 [published Online First: 20230810].

2. Bowers RM, Clum A, Tice H, et al. Impact of library preparation protocols and template quantity on the metagenomic reconstruction of a mock microbial community. BMC Genomics 2015;16:856 doi: 10.1186/s12864-015-2063-6 [published Online First: 20151024].

3. Wang C, Zhang L, Jiang X, Ma W, Geng H, Wang X, Li M. Toward efficient and high-fidelity metagenomic data from sub-nanogram DNA: evaluation of library preparation and decontamination methods. BMC Biol 2022;20(1):225 doi: 10.1186/s12915-022-01418-9 [published Online First: 20221008].

4. Chen Y, Murrell JC. When metagenomics meets stable-isotope probing: progress and perspectives. Trends in Microbiology 2010;18(4):157–63 doi: 10.1016/j.tim.2010.02.002.

5. Mojarro A, Hachey J, Bailey R, et al. Nucleic Acid Extraction and Sequencing from Low-Biomass Synthetic Mars Analog Soils for In Situ Life Detection. Astrobiology 2019;19(9):1139–52 doi: 10.1089/ast.2018.1929 [published Online First: 20190611].

6. Jouvenot Y, Obert C, Hale B, et al. The use of iconPCR for 16S library preparation improves data quality and workflow. bioRxiv 2024:2024.12. 18.629279 doi: 10.1101/2024.12.18.629279.

7. Lu J, Rincon N, Wood DE, et al. Metagenome analysis using the Kraken software suite. Nat Protoc 2022;17(12):2815–39 doi: 10.1038/s41596-022-00738-y [published Online First: 20220928].

8. Beghini F, McIver LJ, Blanco-Miguez A, et al. Integrating taxonomic, functional, and strain-level profiling of diverse microbial communities with bioBakery 3. Elife 2021;10 doi: 10.7554/eLife.65088 [published Online First: 20210504].

9. Green SJ, Venkatramanan R, Naqib A. Deconstructing the polymerase chain reaction: understanding and correcting bias associated with primer degeneracies and primer-template mismatches. PLoS One 2015;10(5):e0128122 doi: 10.1371/journal.pone.0128122 [published Online First: 20150521].

10. Kahsen J, Sherwani SK, Naqib A, Jeon T, Wu LYA, Green SJ. Quantitating primer-template interactions using deconstructed PCR. PeerJ 2024;12:e17787 doi: 10.7717/peerj.17787 [published Online First: 20240808].

11. Bray JR, Curtis JT. An ordination of the upland forest communities of southern Wisconsin. Ecological monographs 1957;27(4):326–49 doi: 10.2307/1942268.

12. Team RC. R: A language and environment for statistical computing. R foundation for statistical computing, Vienna, Austria 2021 doi: http://www.R-project.org.

13. Wickham H, Averick M, Bryan J, et al. Welcome to the Tidyverse. Journal of open source software 2019;4(43):1686 doi: 10.21105/joss.01686.

14. Villanueva RAM, Chen ZJ. ggplot2: elegant graphics for data analysis: Taylor & Francis, 2019.

15. Pedersen TL. Package ‘patchwork’. R package http://CRAN. R-project. org/package= patchwork. Cran 2019 doi: https://CRAN.R-project.org/package=patchwork.

16. Sbaghdi T, Jagorel F, Monot M, Garneau JR. Short-read and Long-read PCR-Free Sequencing of Bacteriophages Using Ultra-Low Starting DNA Input. J Biomol Tech 2025;36(1) doi: 10.7171/3fc1f5fe.c0001573 [published Online First: 20250331].

17. Rinke C, Low S, Woodcroft BJ, et al. Validation of picogram- and femtogram-input DNA libraries for microscale metagenomics. PeerJ 2016;4:e2486 doi: 10.7717/peerj.2486 [published Online First: 20160922].

18. Ospino MC, Engel K, Ruiz-Navas S, Binns WJ, Doxey AC, Neufeld JD. Evaluation of multiple displacement amplification for metagenomic analysis of low biomass samples. ISME Commun 2024;4(1):ycae024 doi: 10.1093/ismeco/ycae024 [published Online First: 20240212].

19. Probst AJ, Weinmaier T, DeSantis TZ, Santo Domingo JW, Ashbolt N. New perspectives on microbial community distortion after whole-genome amplification. PLoS One 2015;10(5):e0124158 doi: 10.1371/journal.pone.0124158 [published Online First: 20150526].

20. Tighe SW, Vellone DL, Tracy KM, Lynch DB, Finstad KH, McLlelan MC, Dragon JA. Microbiome and Microbial Profiling of Arctic Snow Using Whole Genome Sequencing, Psychrophilic Culturing, and Novel Sampling Techniques. J Biomol Tech 2025;36(1) doi: 10.7171/3fc1f5fe.0f37be73 [published Online First: 20250324].

